# Evolutionary increases in catecholamine signaling may underlie the emergence of adaptive traits and behaviors in the blind cavefish, *Astyanax mexicanus*

**DOI:** 10.1101/724179

**Authors:** Kathryn Gallman, Daihana Rivera, Daphne Soares

## Abstract

Evolutionary changes in catecholamine neurotransmitters such as dopamine and noradrenaline can lead to habitat specific behaviors. We used tyrosine hydroxylase, a conserved precursor to the biosynthesis of dopamine and noradrenaline, to compare catecholaminergic neurons in the brain of a species undergoing allopatric speciation. The teleost fish *Astyanax mexicanus* is extant in two readily available forms, an ancestral river dwelling form (surface) and various derived blind cave forms (cavefish). Adaptation to nutrient poor cave life without predation has led to marked differences in the behavior of this species. The cavefish has lost defensive responses, such as stimulus aversion, found in the ancestral surface fish and instead displays enhanced food seeking behaviors. This is reflected by an increase in catecholamine immunoreactivity in the cavefish brain in regions associated with non-visual sensory perception, motor control pathways, attention, and endocrine release. These neuroanatomical regions include the olfactory system, the basal telencephalon, the preoptic nuclei, the posterior tuberculum, caudal hypothalamus, and isthmus. These results indicate that the evolutionary shift from aversive defensive responses to attractive exploratory behaviors was driven by increases in the size and/or quantity of catecholaminergic neurons in the cavefish brain.

## Introduction

Changes in neuromodulation can facilitate the emergence of new behaviors as a diverging species adapts to a novel habitat. One of the most common classes of neuromodulators in the central nervous system are the catecholamines, which include dopamine (DA) and noradrenaline (NA). Catecholamines mediate a wide array of behaviors, such as, cognition, memory, emotion, temperature regulation, sleep, and motor control (Björklund & Dunnett, 2000). Although commonly considered in the context of reward or aversive stimuli association, catecholamines primarily select the action, or response behavior, generated by a particular stimulus (Chakravarthy et al., 2010).

Catecholamine signaling selects for survival seeking behaviors, such as foraging for food. This “seeking” system (for review see Wright & Panksepp, 2014) can be activated through both internal homeostatic signals that indicate states like hunger and external sensory stimuli that indicate changes in the environment, like the appearance of a nutrient source. Stimuli that detect food and behaviors that successfully acquire food are reinforced through catecholaminergic potentiation of sensory and motor circuit synapses (Schultz, 1998; Calabresi et al., 2007). NA signaling acts as an attention filter to highlight crucial or novel stimuli (for review see Prokopova, 2010), while DA transmission increases the rate of the locomotor response (Purves, 2008). This allows an animal to both anticipate and respond more quickly to similar future stimuli (Chakravathy et al., 2010).

Variations in DA and NA neurons between species may account for instinctual behaviors that shaped adaptation. Tyrosine hydroxylase (TH) is the rate limiting enzyme in the biosynthesis of DA and NA and highly conserved across taxa (Candy & Collet, 2005; Wall & Volkoff, 2013). By examining differences in TH immunoreactivity (THir) in the brains of a species undergoing allopatric speciation, we hope to better understand the neural correlates of behavioral adaptation to novel environments. Here we compare surface and cave dwelling forms of the teleost fish *Astyanax mexicanus*, because the evolution to cave life has created a unique set of survival challenges. Unlike the dynamic river waters, water in the Pachón cave of the Sierra del Abra region of Mexico is relatively stagnant for most of the year and receives little nutrient influx (Rétaux et al., 2015). At the top of the cave food chain, the primary threat to the survival of the cavefish is starvation (Trajano et al., 2010). This has created a runaway effect on food seeking behaviors (Salin et al., 2010; Duboue et al., 2011; Rétaux et al., 2015). To this effect, we discovered increases in the size and/or quantity of THir neurons in regions of the cavefish brain associated with detecting and finding food in the darkness.

## Methods

### Animals

Adult specimens of surface and Pachón cave *Astyanax* (standard lengths of 4.7-5.1 cm, n=20) were raised under a 12-hour light-dark cycle and fed once daily. All experimental procedures were approved by the Institutional Animal Care and Use Committee of Rutgers University Newark (protocol: 201702685).

### Tissue Preparation

Fish were euthanized via immersion in overdosed pharmaceutical grade tricaine methanesulfonate salt (MS-222, 10g/L), and perfused transcardially with cold heparinized saline (0.9% Sodium Chloride with heparin (20 units/mL) followed by fixation in 10% neutral buffered formalin (4% formaldehyde). Fish were decapitated and heads post fixed for 4 hours before transfer to a 30% sucrose solution overnight at 4°C. Brains were removed from the skull via dissection the following day and further cryoprotected in 30% sucrose in 4°C prior to sectioning. Serial sectioning was performed at −20°C (cryostat: ThermoFisher HM525NX) in coronal, sagittal, and horizontal planes and mounted on positively charged glass slides. Sections were either processed the following day or stored at −20°C until one day prior to processing.

### Immunohistochemistry

Slide mounted sections were rehydrated in 0.1M PBS twice for ten minutes, washed, and then blocked for 2 hours at room temperature (0.2% fetal bovine serum (FBS), 0.3% Tween 20, 10% normal goat serum (NGS) in 0.1M PBS). Sections were incubated overnight at 4°C in primary antibody (TH monoclonal mouse antibody, cat. #: MAB318, lot #: 3083054, Millipore) diluted 1:500 in incubation buffer (0.2% FBS, 0.3% Tween 20 in 0.1M PBS). The following morning, sections were washed thrice for 10 minutes with 0.1M PBS and 0.1% Tween20 (PBSTween) and incubated with Alexa Fluor 594 conjugated anti-mouse secondary antibody (abcam, ab150116) diluted 1:500 in blocking solution for two hours at room temperature. This was followed once more by three 10-minute washes with PBSTween. Slides were then counterstained with DAPI (1:5000) for 30 minutes and washed thrice with 0.1M PBS before mounting with 90% glycerol in phosphate buffer.

### Imaging

Immunofluorescent images were taken using a confocal microscope (Leica SP8) with excitation at 405 nm and 552 nm laser lines. Scans of full sections were acquired with 10x and 20x dry objective lenses and Leica LAS X Navigator software. Scans of THir nuclei were taken with 40x and 63x oil immersion objective lenses. Images were post processed using Leica LAS X confocal software.

### Quantification

20 μm (average cell body size) coronal serial sections were imaged using an epifluorescent microscope (Olympus PX50) at 40x magnification with an OMAX 18.0MP digital microscope camera and Toupe software. Cell bodies diameters were measured from the axon hillock to the farthest point of the soma using FIJI (Schindelin et al., 2012). Log transformed cell sizes by nuclei were then compared between surface and cave *Astyanax* of the same standard length using a student’s t-test in RStudio.

### Nomenclature

Because an anatomical atlas of *Astyanax* is not yet available, nuclei identification was based on work in other teleost fishes including: Wullimann et al. (1996), Rincón et al (2017), and Biechl et al. (2017) with updated periventricular organ (PVO) and posterior tuberal nucleus (PTN) boundaries defined in Rink and Wullimann (2001).

## Results

### General observations

THir nuclei were found primarily in the telencephalon and diencephalon. Rostrally to caudally these regions included: the olfactory bulb (OB), the ventral telencephalon, the preoptic area, the ventral thalamic area, posterior tuberculum, and the caudal hypothalamus. There were no THir cell bodies in the mesencephalon, although there was a group of large THir cell bodies in the isthmus region, or the boundary between the mesencephalon and rhombencephalon. There were also THir neurons in the area postrema and vagal lobe of the telencephalon of *Astyanax* (not shown).

THir fibers were most prominent within and projecting from brain nuclei with THir cell bodies, but also in rhombencephalon nuclei associated with gustatory, auditory, and lateral line processing. Overall, the density of cells and fiber projections appears greater in the cave *Astyanax* than its surface counterpart, particularly with respect to olfactory projections and the nuclei of the posterior tuberculum.

### Olfactory bulb

The OB of the cavefish was larger than the surface fish with more interneurons (Menuet et al., 2017). The overall shape of this structure also differed between morphotypes. In cross sections, the surface fish OB was almost triangular in shape with the rounded apexes of the left and right OB angled towards each other, while the cavefish OB was more ellipsoid in shape with the rounded apexes angled away from each other. These differences in shape were also reflected by differences in THir cell bodies and their processes (Fig. 1).

**Figure 1:**
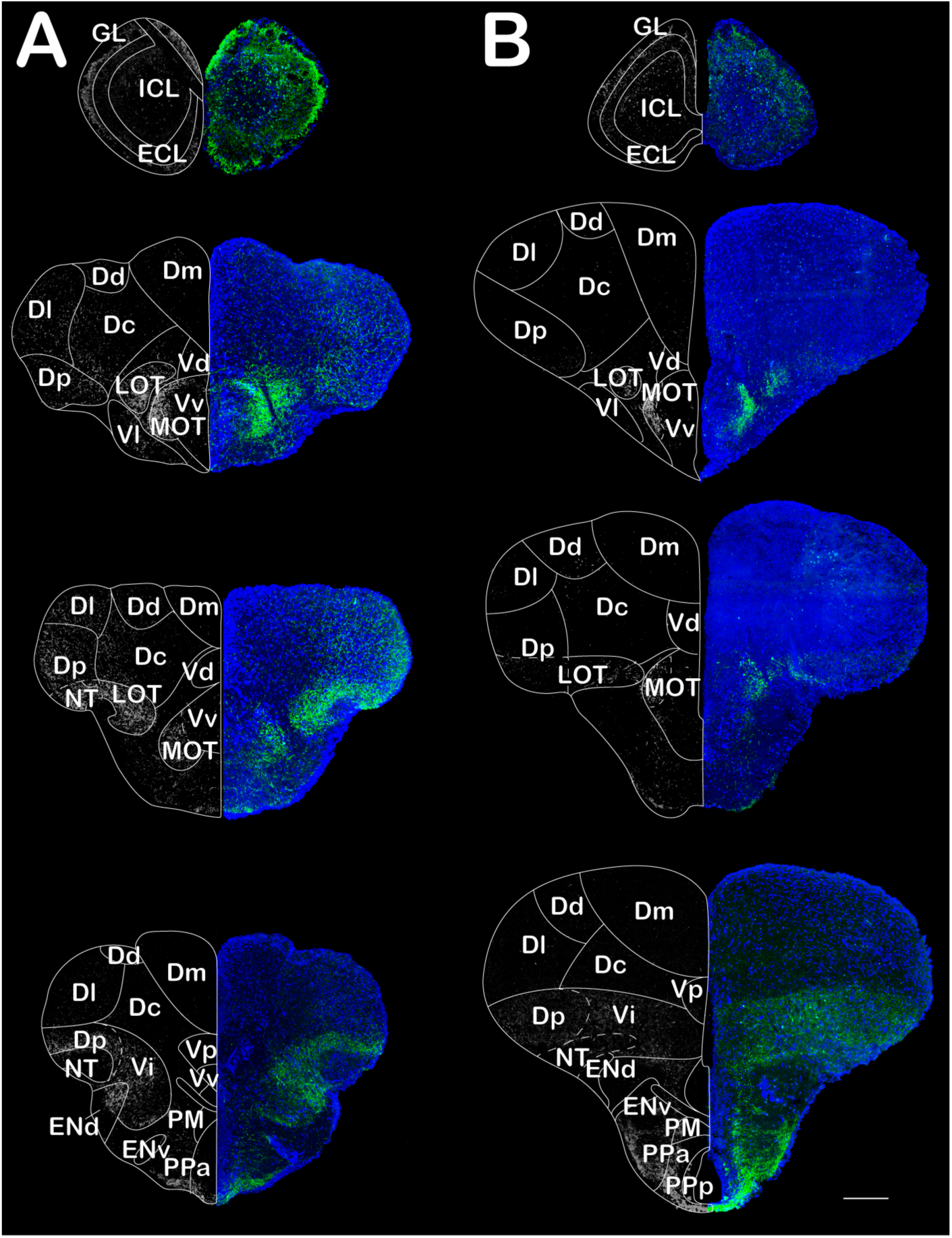
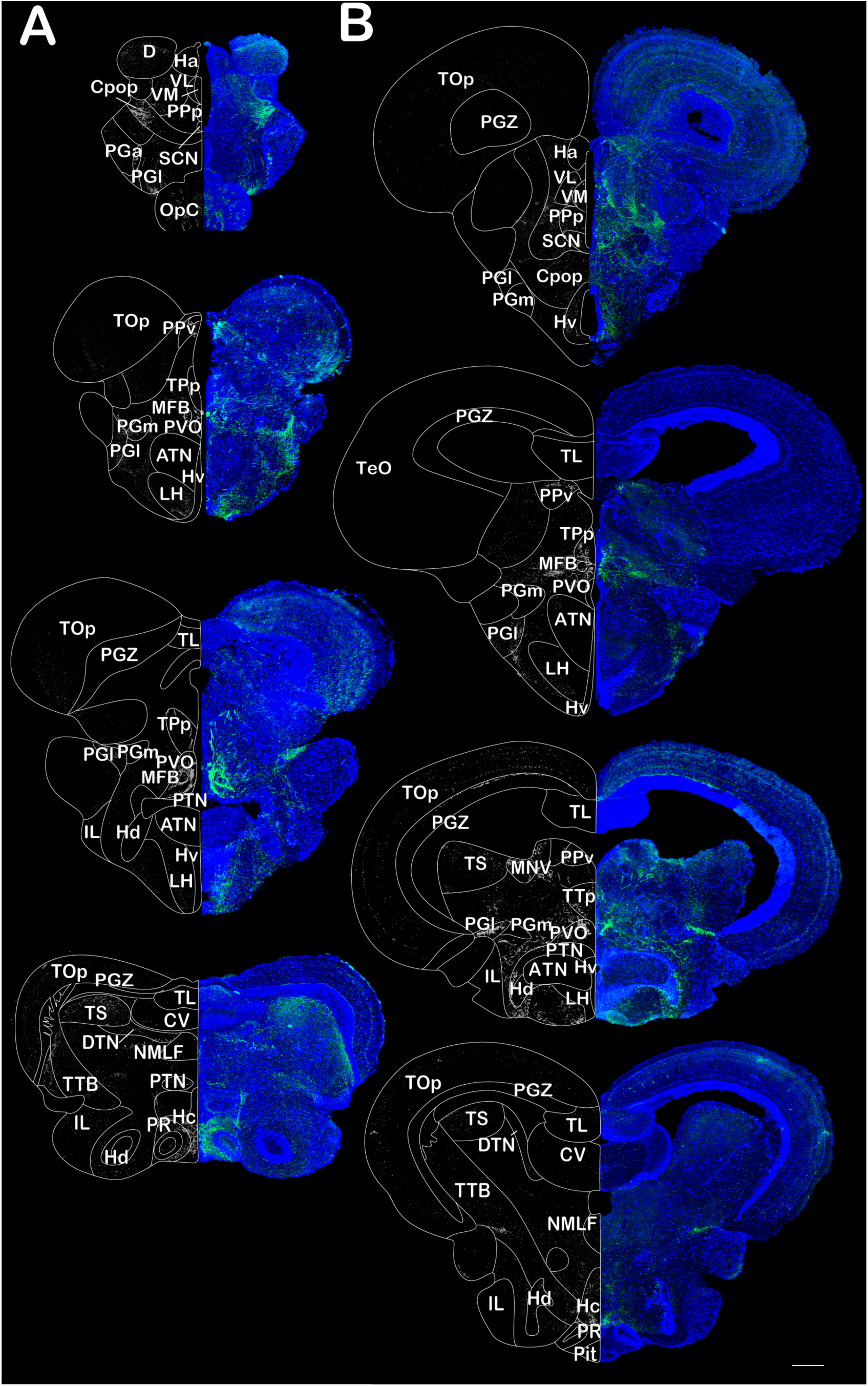
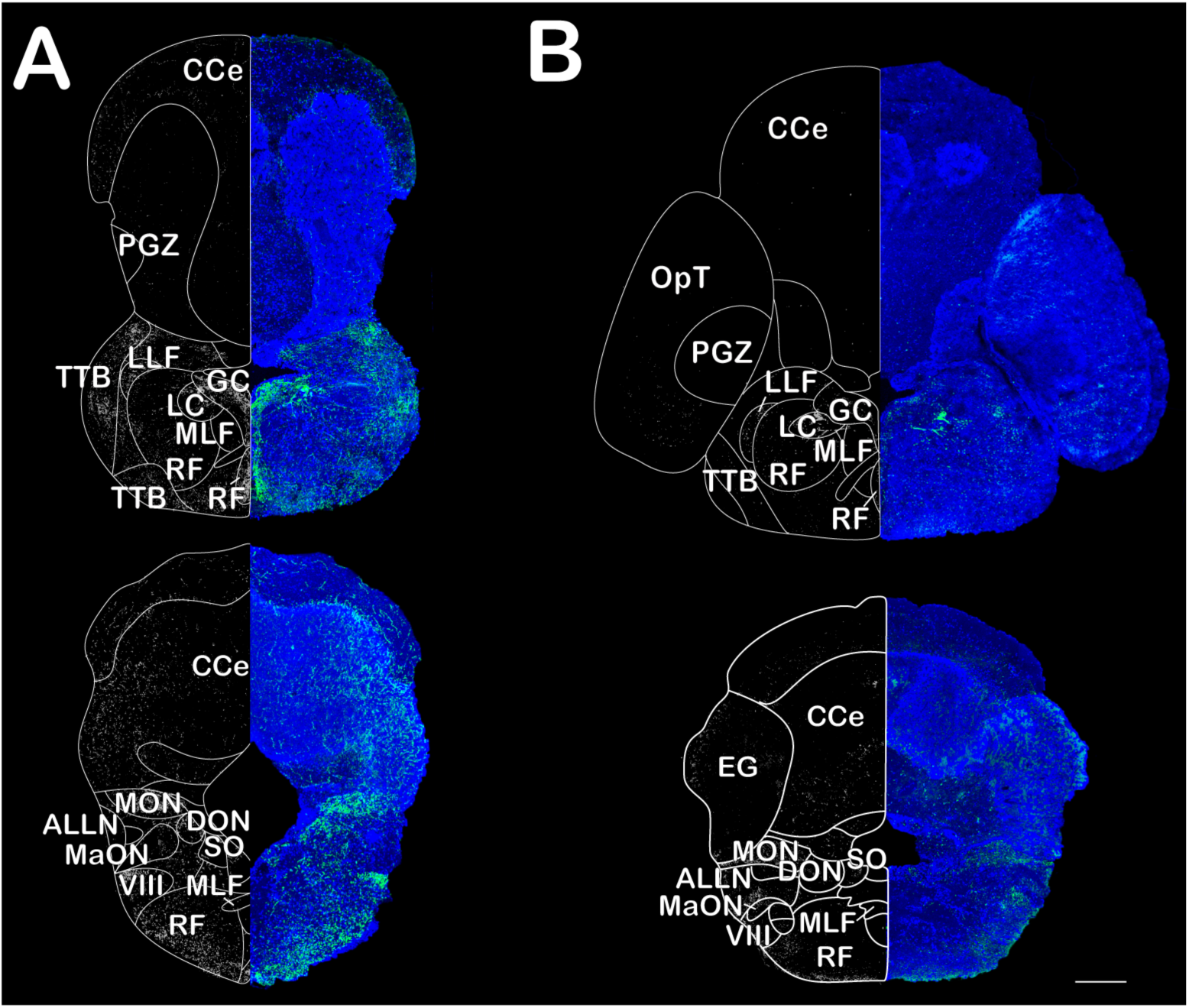
Comparative THir (green/white) Astyanax confocal coronal sections counterstained with DAPI (blue) between cavefish (column A) and surface fish (column B). Columns run rostral to caudal. Scale bars are 80 μm.

In both fish, THir soma were located predominantly along the border of the inner cell layer (ICL) and the external cell layer (ECL) and sent projections to the dendrites of mitral cells in glomerular layer (GL) in clusters. Surface fish THir cells were significantly larger (p-value < 0.001) than those of the cavefish with an average cell size of 12.5 ± 2.1 μm, although there were fewer cells overall in the surface fish ICL (n = 2,831 cells) compared to the cavefish (n = 3,252 cells) and fiber projections were limited to the lateral and dorsal regions of the GL (Fig. 2). The average cell size for cavefish THir neurons (6.7 ± 1.2 μm) was approximately half that of the surface fish, but these smaller cells appeared greater in density and with many more fiber projections innervating almost the entire GL.

**Figure 2:**
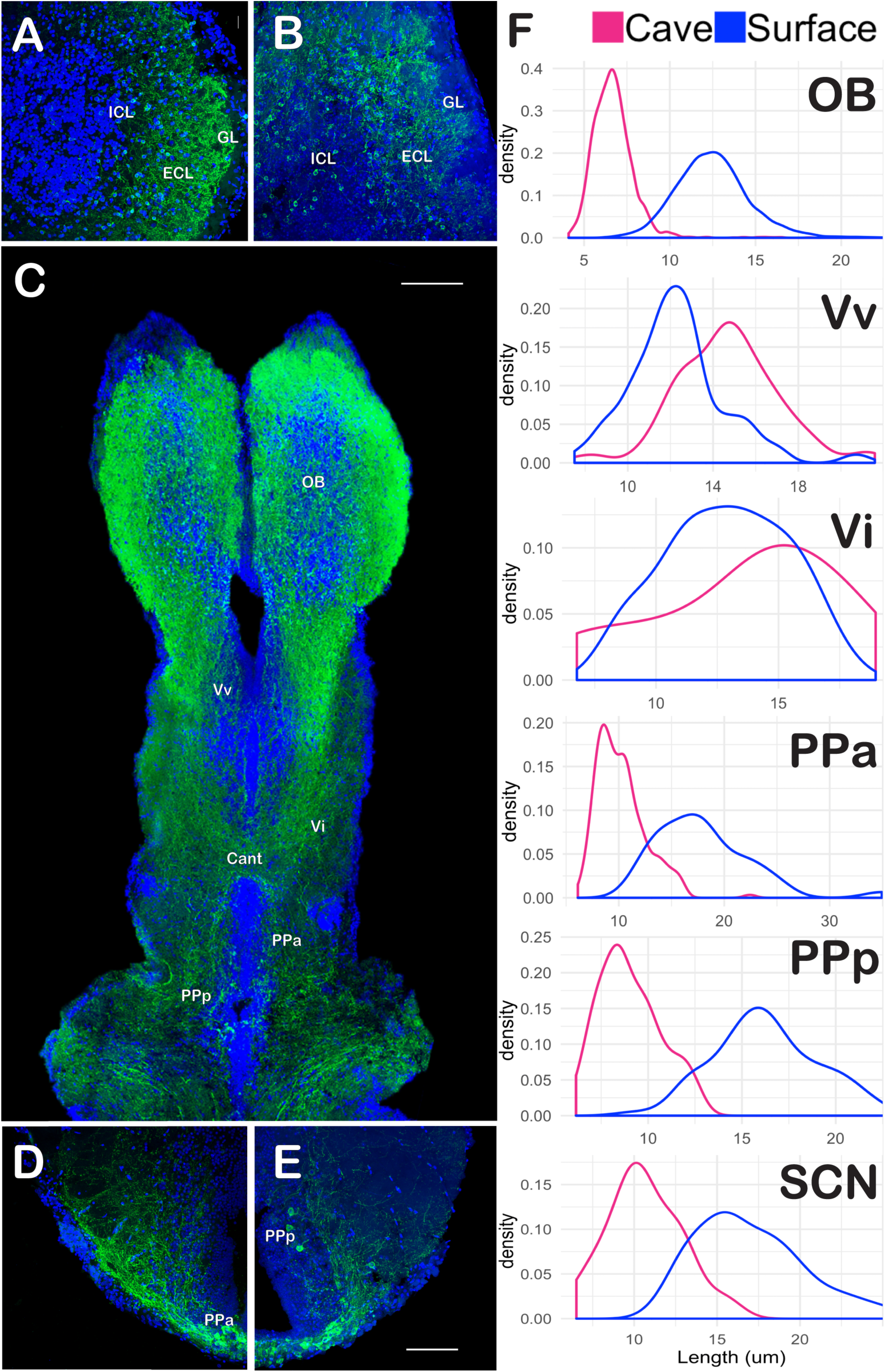
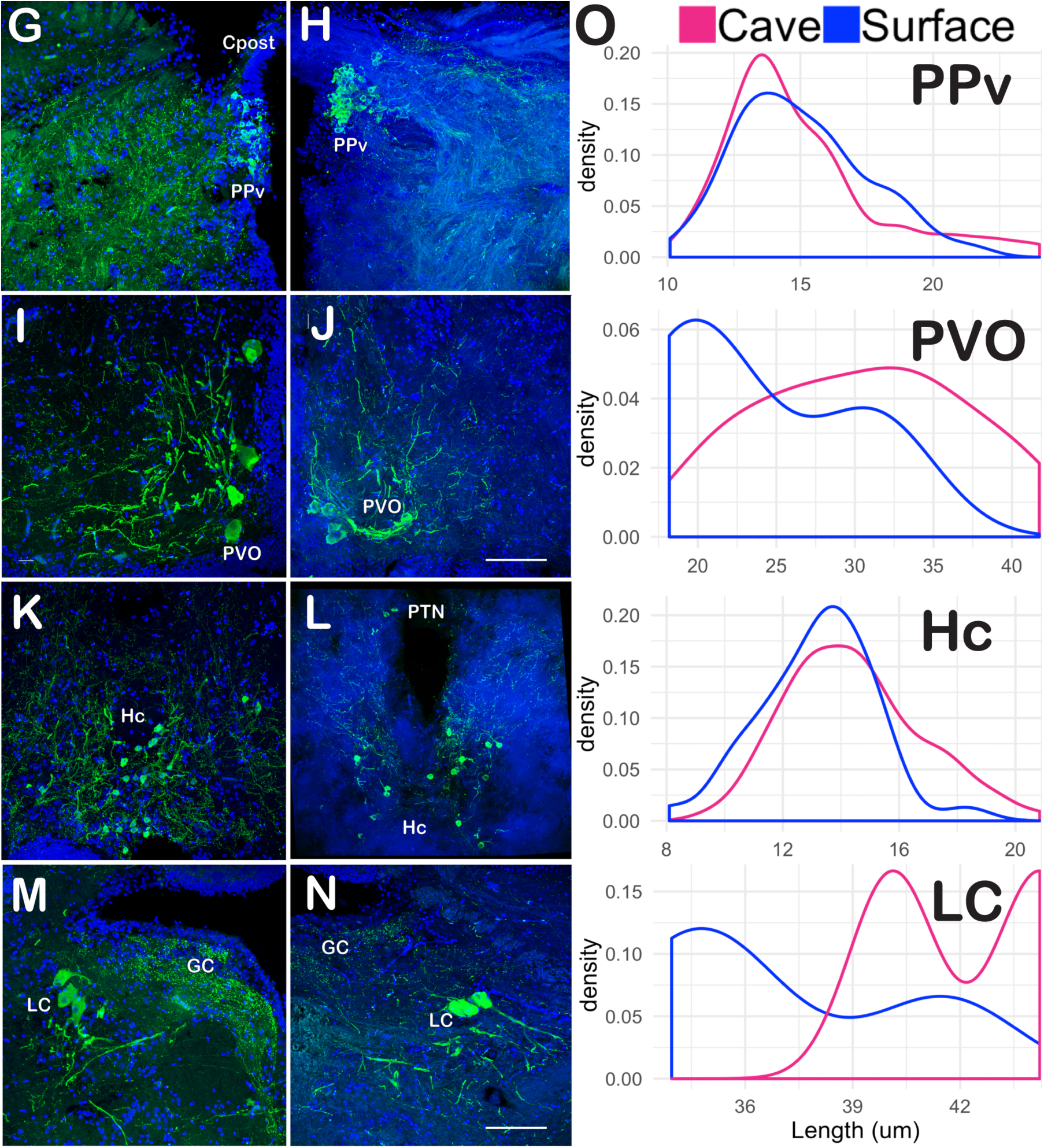
In the olfactory bulb (OB), anterior (PPa) and posterior (PPp) parvocellular preoptic nuclei, and suprachiasmatic nucleus (SCN), the cavefish has more THir cell bodies of a significantly smaller diameter than in the surface fish (p-value < 0.001). In the ventral (Vv) and intermediate (Vi) zones of the ventral pallium, the frequency of THir cell bodies was similar between cave and surface fish, although cavefish had significantly larger Vv cell body diameters (p-value < 0.001). **A-E)** Confocal scans of THir nuclei (green) counterstained with DAPI (blue). **A, B)** Coronal sectioned OB of cave (A) and surface (B) *Astyanax*. **C)** Horizontal sectioned *Astyanax* cavefish brain showing olfactory projections from the OB through the telencephalon at the level of the anterior commissure (Cant). Scale bar 500 μm. **D, E)** Coronal sectioned basal telencephalon of cave (D) and surface (E) *Astyanax* imaged at 40x magnification. Scale bar for A, B, D, and E 80 μm. **F)** Kernel density estimates (smoothed histograms) of cell body diameter measurements in micrometers. THir cell bodies of the Posterior periventricular nucleus (PPv) varied little in number and diameter between surface and cave *Astyanax*. There were very few THir cell bodies in the Paraventricular organ (PVO) and the locus coeruleus (LC), but these neurons were larger and more numerous in the cavefish. The cavefish also had significantly more cell bodies of larger size in the caudal hypothalamus (Hc; p-value < 0.01). **G-N)** Coronal sectioned confocal scans of THir nuclei (green) counterstained with DAPI (blue). The left column (G, I K, M) shows scans of the cavefish brain, while the right column (H, J, L, N) shows scans of the surface fish brain. All scale bars are 80 μm. **G, H)** THir PPv cells occurred in dense clusters just ventral to the posterior commissure (Cpost). **I, J)** THir cell bodies of the PVO were large and pear shaped (Type 2 neurons) and projected in a semicircle from the diencephalic ventricle (DiV). **K, L)** Hc THir cells appeared most densely in the caudal and ventral portion of the nucleus, clustered around the DiV. **M, N).** LC THir cells bodies had large diameters and occurred just ventral and lateral to the griseum centrale (GC). **O).** Kernel density estimates (smoothed histograms) of cell body diameter measurements in micrometers.

### Telencephalon

Like the OB, the gross morphology of the telencephalon also differed in shape between surface and cave *Astyanax*. The surface fish had a much larger dorsal pallium while the cavefish telencephalon appeared much smaller, but with a broader ventral pallium. This contrast in overall morphology influenced differences in the shape of mitral cell axon projections (medial and lateral olfactory tracts) as they moved caudally through the telencephalic region (Fig. 1).

THir medial olfactory tract (MOT) cells and fibers appeared in the ventral region of the ventral pallium (Vv) and the lateral olfactory tract (LOT) fibers appeared to send the majority of projections to the dorsal region of the dorsal pallium (Dp), although these extended much further dorsally and laterally with greater complexity in the cavefish. The fiber projections of the LOT were immediately lateral to those of the MOT and the two tracts could be differentiated from one another visually by a gap in THir fibers between the populations (Fig 1).

The shape of the transition from MOT to LOT, in the coronal plane, reflected the differences in *Astyanax* telencephalon morphology. In the cavefish, olfactory fibers extended laterally from Vv towards Dp. In the surface fish, these fibers first had to move dorsally from Vv to Vd before they were able to travel laterally towards Dp, and as a result, had a small population of THir cells in Vd that are absent in the cavefish (Fig. 1).

Although immunoreactivity for Vv occurred mainly on the lateral border of the nucleus rather than medially near the ventricle, we have classified these cell bodies and their processes as the MOT termination within Vv, as this more lateral location was consistent with reports of THir in other teleosts. The density of cell bodies in Vv did not widely vary between surface (n=119 cells) and cave *Astyanax* (n=128 cells); however, the cell bodies of the cavefish were significantly larger (p-value < 0.001) than those of the surface fish with an average of 15 ± 2.4 μm compared to 12 ± 2.3 μm (Fig 2). The dense THir fibers of the cavefish also covered a much broader area of the subpallium than those of the surface fish.

The surface fish also showed THir fiber projections to the dorsal region of the dorsal pallium (Dd) that decreased in density dorsally (Fig 1). This region was unique among telencephalic THir in that the immunoreactivity did not occur in the cytoplasm but in small puncta clustered around the cell bodies (not shown). This staining pattern and location of THir was completely absent in the cavefish.

Caudal to Vv, there was a second set of cells and fibers along the pallial-subpallial boundary that we have attributed to the intermediate zone of the ventral pallium (Vi). In this part of the telencephalon, the olfactory projections converged to cross the midline through the telencephalic commissure and continued their projections through Vi to the dorsal pallium, weaving around the lateral forebrain bundles (LFB; Fig. 1). The cell bodies of this region averaged 12.9 ± 2.5 μm for the surface fish and 13.6 ± 3.9 μm for the cavefish with no significant size differences (Fig. 2) but are scattered amongst the LFB more densely in the surface fish (SF n= 51 cells compared to CF n=14 cells).

Vi fibers increased in density caudally and became intertwined with the dorsally projecting fibers of the THir cell bodies of the anterior region of the parvocellular preoptic nucleus (PPa; Fig 1). Similar to the OB, there were much more PPa THir cell bodies in the cavefish (n= 411) than the surface fish (n= 36), but although the cavefish PPa cells appeared in greater densities, they are also significantly smaller in size (average of 10.2 ± 3.4 μm compared to 18.0 ± 4.6 μm; Fig 2). The majority of the fiber projections from these regions terminated in Dp, although there were sparse projections throughout most of the dorsal pallium.

### Abbreviations

ALLN: Anterior Lateral Line Nerves
ATN: Anterior Tuberal Nucleus
Cant: Anterior commisure
Cpost: Posterior commisure
CCe: Corpus cerebelli
CV: Valcula cerebelli
D: Dorsal pallium
Dc: Central zone of D
Dd: Dorsal zone of D
Dl: Lateral zone of D
Dm: Medial zone of D
Dp: Posterior zone of D
DTN: Dorsal tegmental nucleus
ECL: External cell layer
EG: Eminentia granularis
ENd: Dorsal Entopeduncular nucleus
Env: Ventral Entopeduncular nucleus
GC: Griseum centrale
GL: Glomerular layer
Ha: Habenula
Hc: Caudal zone of periventricular hypothalamus
Hd: Dorsal zone of periventricular hypothalamus
Hv: Ventral zone of periventricular hypothalamus
ICL: Internal cell layer
IL: Inferior lobe of hypothalamus
LC: Locus coeruleus
LH: Lateral hypothalamus
LLF: Lateral longitudinal fasicle
LOT: Lateral olfactory tract
MaON: Magnocellular octaval nucleus
MFB: Median fiber bundle
MLF: Medial longitudinal fasicle
MON: Medial octavolateralis nucleus
MOT: Medial olfactory tract
NMLF: Nucleus of the medial longitudinal fasicle
PGl: Lateral preglomerular nucleus
PGm: Medial preglomerular nucleus
PGZ: Periventricular gray zone of the optic tectum
PM: Magnocellular preoptic nucleus
Ppa: Parvocellular preoptic nucleus
PPp: Posterior part of the parvocellular preoptic nucleus
PPv: Posterior periventricular nucleus
PR: Posterior recess of the diencephalic ventricle
PTN: Posterior tuberal nucleus
PVO: Paraventricular organ
RF: Reticular formation
SCN: Suprachiasmatic nucleus
SGN: Secondary gustatory nucleus
SO: Secondary octaval population
TOp: Optic tectum
TPp: Periventricular nucleus of the posterior tuberculum
TPN: Thalamic posterior nucleus
TS: Torus semicircularis
TTB: Tecto-bulbar tract
V: Ventral telencephalic area
Vd: Dorsal nucleus of V
Vi: Intermediate subdividsion of V
VIII: Octaval nerve
Vl: Lateral nucleus of V
VL: Ventrolateral thalamic nucleus
VM: Ventromedial thalamic nucleus
Vp: Postcomissural nucleus of V
Vv: Ventral nucleus of V

### Preoptic area

In sagittal view, PPa could be seen as a line of THir neurons with dorsally projecting axons that ran along the base of the telencephalon. At the transition to the diencephalon, this line became curved as the neurons of the posterior region (PPp) climbed the dorsal edge of the optic tract (Fig. 5). The distinction between parvocellular nuclear regions was necessary because PPp has significantly more neurons of smaller size than PPa for both *Astyanax* morphotypes (p-value < 0.05). The PPp region of the cavefish was also significantly smaller than in the surface fish (p-value <0.001) with a mean of 9.0 ± 1.6 um compared to 16.3 ± 2.8 (Fig. 2). There also appeared to be fewer total PPp THir neurons in the cavefish (n=66 cells) than in the surface fish (n=112 cells)

**Figure 3:**
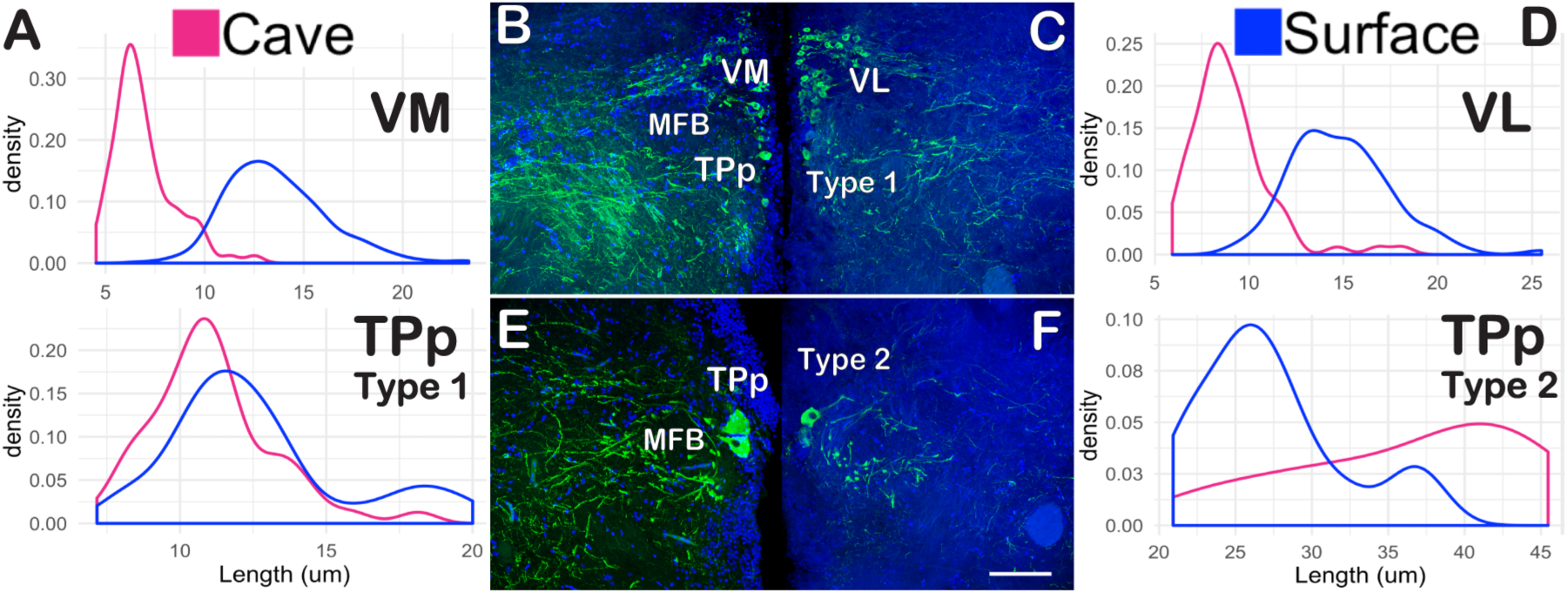
THir cell bodies in the ventromedial (VM) and ventrolateral (VL) thalamus were significantly larger in the surface fish brain than in the cavefish (p-value < 0.001). THir Type 1 cell bodies in the periventricular nucleus of the posterior tuberculum (TPp) were also significantly larger in the surface fish than in the cavefish (p-value < 0.01). Although there were only a few THir Type 2 TPp cell bodies, these appeared larger and more numerous in the cavefish. **B, C, E, F)** Coronal sectioned *Astyanax* confocal images of THir nuclei (green) counterstained with DAPI (blue). The left column (B, E) shows the cavefish brain, while the left column (C, F) shows the surface fish brain. All scale bars are 80 μm. **A, D)** Kernel density estimates (smoothed histograms) of cell body diameter measurements in micrometers.

**Figure 4:**
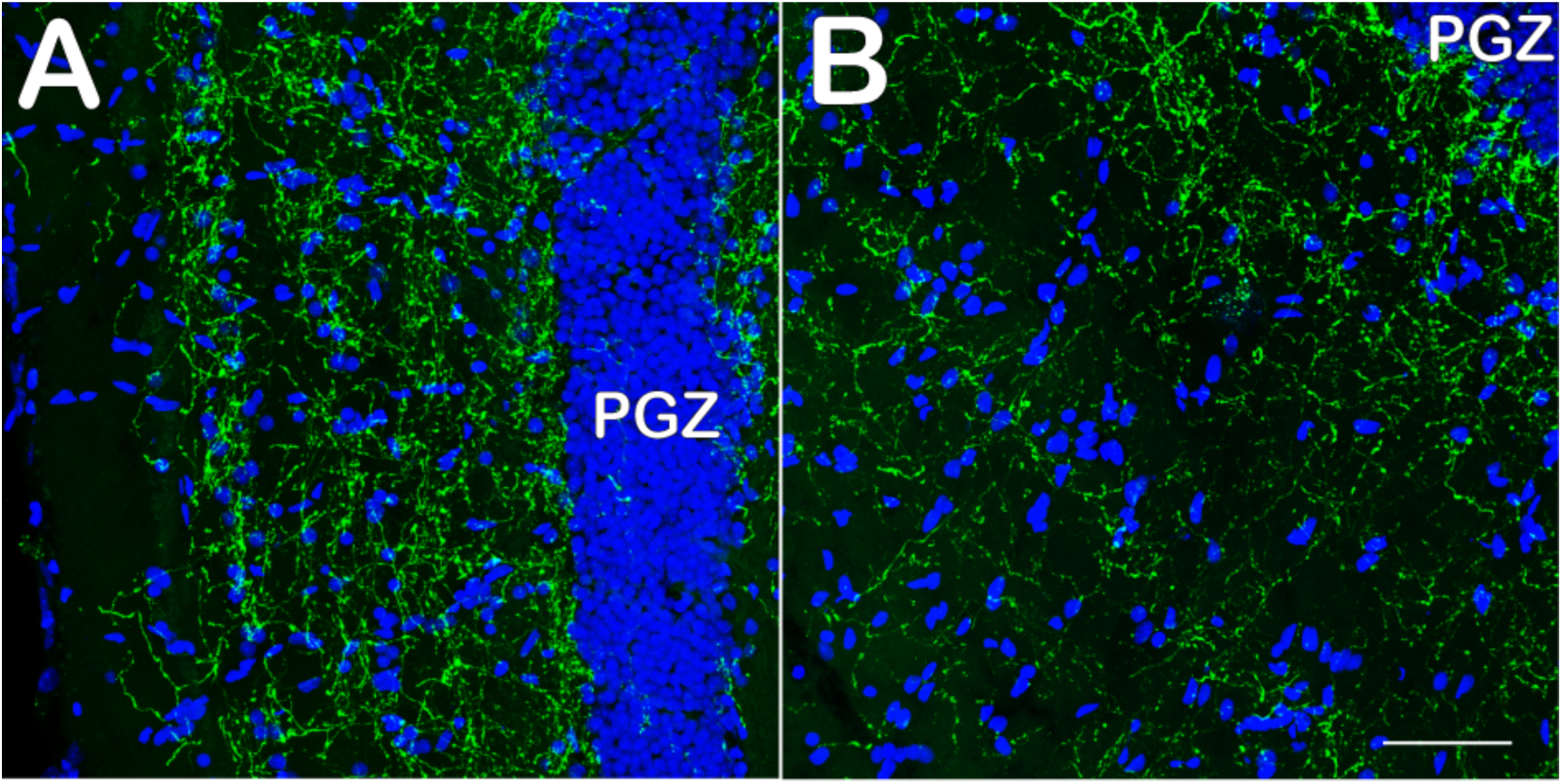
THir appears in the optic tectum of cave (A) and surface (B) *Astyanax* in layered bands. Confocal scans of horizontal sections immunolabeled with TH (green) and DAPI (blue). Scale bar 80 μm.

**Figure 5:**
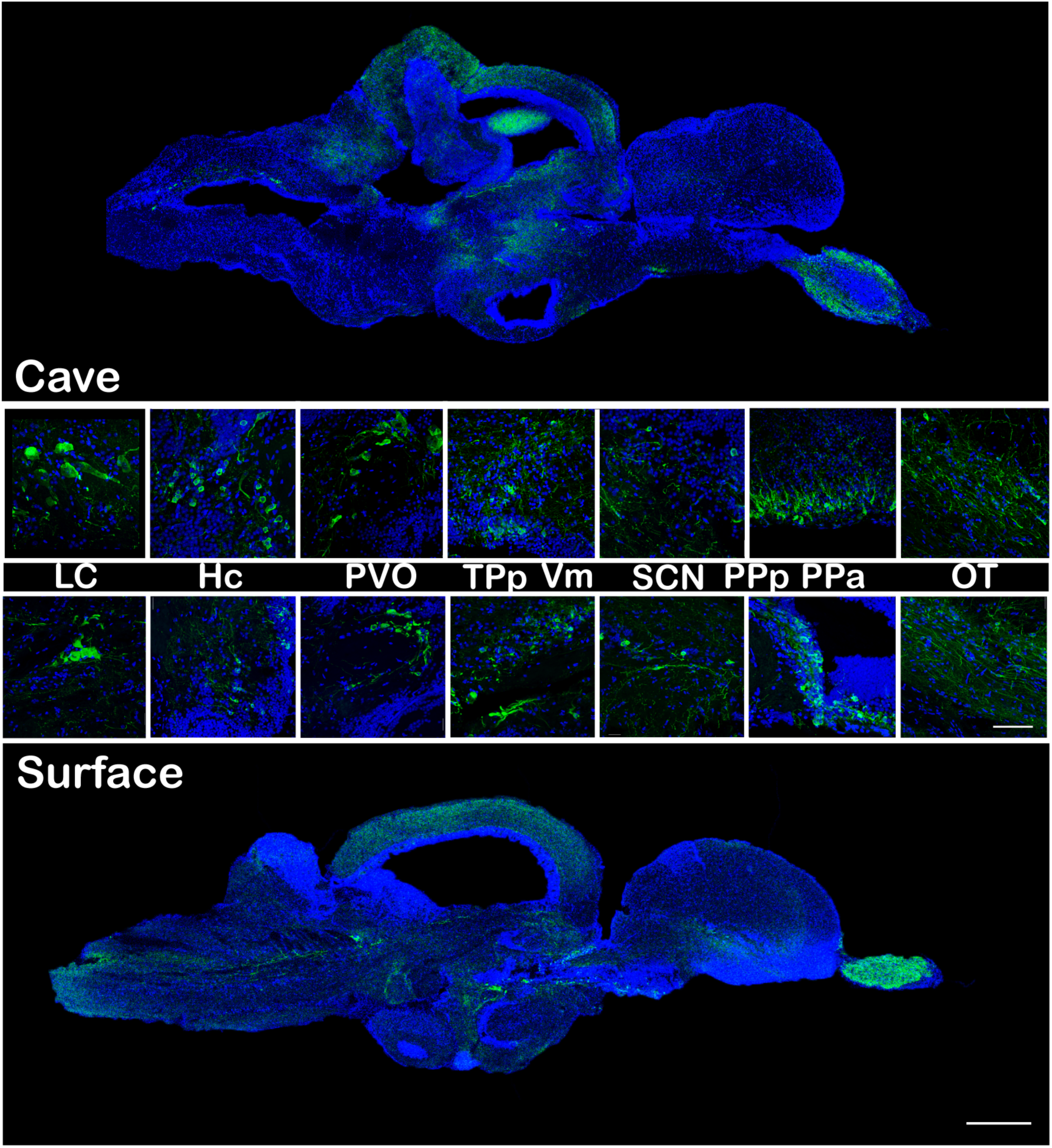
Cave (top) and surface (bottom) *Astyanax* THir in the sagittal plane, 500μm scale bar. Inserts 80 μm scale bar. Immunolabeled for TH (green) and DAPI (blue).

The optic tract crossed just caudal of PPp and immediately dorsal to this chiasm, the suprachiasmatic nucleus (SCN) also exhibited TH immunoreactivity (Fig. 1). The surface fish SCN had significantly more of large cell bodies (p-value < 0.001; n=38 cells) than the cavefish (n=17 cells) with a mean of 17.0 ± 3.2 compared to 10.6 ± 2.2 (Fig. 2).

### Thalamus

PPp and SCN appeared ventral to the habenular region in coronal section. Although THir projections to the habenular region were sparse, dense populations of ventral thalamic neurons began just dorsal to the habenula and extend ventrocaudally to the tuberal region (Fig. 5). Cell bodies of the ventromedial thalamic nucleus (VM) were located along the ventricle and could be distinguished from cell bodies of the ventrolateral thalamic nucleus (VL) both by their more lateral location and by a significant increase in size. The average diamter of the cavefish Vm cell bodies was 7.0 ± 1.5 μm compared to 9.0 ± 2.1 μm for VL cell bodies (p-value < 0.001). The average diameter of the surface fish VM was 13.6 ± 2.6 μm compared to 14.9 ± 2.7 μm for VL cell bodies (p-value < 0.001). In the coronal plane, lateral projections from both regions coalesced into a distinctive band that tapered off horizontally, at a slightly downward angle, across the coronal plane. Cell bodies of VM and VL appeared more numerous (SF n= 295 cells compared to CF n=77 cells for VM; SF n= 96 cells compared to CF n=78 cells for VL) and were significantly larger (p-value < 0.001) in the surface fish than the cavefish with means of 13.7 ± 2.6 μm and 70 ± 1.5 μm for VM and 14.9 ± 2.7 μm and 90 ± 2.1 μm for VL (Fig. 3).

### Synencephalon

There was a densely packed group of THir cells within the periventricular pretectal nucleus (PPv) just ventral to the posterior commissure that sent projections over the fasciculus retroflexus (FR) into the optic tectum (TOp; Fig. 1). THir fibers themselves could be seen in the TOp in distinctive bands in both *Astyanax* forms (Fig. 4). There was no significant difference in cell body size between surface and cave *Astyanax* THir PPv neurons as both morphotypes have a mean of 15 μm with a standard deviation of ± 2.9 μm for the cavefish and ± 2.5 μm for the surface fish (Fig. 2). However, there appeared to be more surface fish (n=228 cells) PPv THir neurons than in the cavefish (n=108 cells).

### Posterior tuberculum

In the sagittal plane, THir ventral thalamic neurons began in the rostral diencephalon and continued ventrocaudally until they reached the periventricular nucleus of the posterior tuberculum (TPp; Fig. 5). TPp THir neurons exhibited a similar ventrolateral projection pattern to the ventral thalamic neurons and these two axon tracts encircled the medial fiber bundle (MFB; Fig. 3). There were at least two separate populations of THir neurons in TPp based on cell body diameter. Following the nomenclature used by Rink & Wullimann (2001), we have designated smaller cell bodies as type 1 neurons (3-20 μm) and larger cell bodies as type 2 (20+ μm) neurons. Although cerebrospinal fluid contacting cells were present, they were not always clearly distinguishable from laterally projecting cells, and therefore, have been included with the smaller type 1 cells for analysis.

Type 1 TPp neurons occurred immediately ventral to the ventral thalamic neurons and were significantly larger (p-value < 0.01) in the surface fish (mean of 12.6 ± 3.1 μm; n=32) compared to the cavefish (mean of 11.1 ± 1.9 μm; n=48). The larger type 2 neurons appeared ventral and caudal to type 1 neurons, with a distinctive pear shape cell body and thicker axons that encircled the MFB (Fig. 3). Type 2 cell bodies only occurred in a few coronal sections and therefore did not provide a large enough sample size for statistical analysis, however, surface fish type 2 neurons ranged mostly from 20 – 30 μm (mean of 27.3 ± 4.8 μm; n=15) while cavefish Type 2 neurons ranged from 35 – 45 μm (mean of 36.1 ± 7.6 μm; n=14), which indicates larger type 2 cell bodies in the cavefish (Fig. 3).

Caudal and ventral to type 2 TPp neurons were another group of large THir cell bodies in the region redefined by Rink and Wullimann (2001) as PVO (Fig. 1). This is important to note because it was previously classified as PTN in THir cell identification in other teleost studies. These cell bodies appeared in a semicircle along the diencephalic ventricle and sent long axonal projections dorso and ventrocaudally. Although, there was not enough power for statistical analysis of PVO neurons, there were visual differences between cave and surface fish PVO neurons. The cavefish cell bodies appeared to be larger, ranging from 20 – 45 μm (mean of 30.5 ± 6.7 μm; n=26), compared to the surface fish that range from 15 – 33 μm (mean of 23.7 ± 6.1 μm; n=6; Fig. 2). Lateral to PVO, in the coronal plane, THir fibers could also be found in the lateral preglomerular nucleus (PGl), but not in the medial preglomerular nucleus for both forms of *Astyanax* (Fig. 1).

Finally, there were only a few small THir cells within the newly defined PTN. These appeared only in the most caudal region of the nucleus and appeared to be associated with the dorsal projections of THir cells in the caudal hypothalamus (Hc; Fig. 2).

### Hypothalamus

The cavefish had a larger hypothalamus than the surface fish (Menuet et al., 2007), which is most apparent in a broadening of the caudal region in the coronal plane. Although fibers appeared in the dorsal (Hd) and lateral hypothalamus (LH), THir cell bodies were only present in Hc (Fig. 1). Hc cell bodies were located medially to the posterior recess and projected dorsolaterally. Although both cave and surface *Astyanax* cell bodies had a mean length of 15.0 μm (with standard deviations of ± 2.9 μm for the cavefish and ± 2.5 μm for the surface fish) the cavefish had significantly more large cell bodies than the surface fish (p < 0.01; SF n= 199 cells compared to CF n=128 cells). There were also THir fibers present in the inferior lobes and pituitary in both *Astyanax* forms (Fig. 2).

### Mesencephalon

Dorsal to Hc, THir fibers were present in torus semicircularis (TS) of the cavefish, but not the surface fish (Fig. 1). Caudal to Hc, THir fibers could be found in the reticular formation (RF; Fig. 1), the lateral longitudinal fascicle (LLF; Fig. 1), the interpeduncular nucleus (IPN; not shown), the tectobulbar tract (TTB; Fig. 1), the secondary gustatory nucleus (SGN, not shown), the thalamic posterior nucleus (TPN, not shown), griseum centrale (GC; Fig. 1, 2), and corpus cerebelli (CC; Fig. 1). There are no THir cell bodies until the locus coeruleus (LC; Fig. 1) in the isthmic region, which displayed a few large cell bodies and projected in the rostral and caudal directions (Fig. 2, 5). These large cell bodies had a mean length of 37.0 ± 4.0 μm in the cavefish and a mean length of 42.2 ± 2.9 μm in the surface fish (Fig. 2). The cavefish LC neurons were more readily discernible than those of the surface fish and occurred more frequently (n = 30 verses n = 15 cell bodies), despite their size similarities.

### Rhombencephalon

Although unquantified, THir fibers appeared to be present in greater densities in the cavefish anterior brainstem than the surface fish, particularly with respect to the secondary gustatory nucleus (SGN; not shown), reticular formation (RF; not shown), and several octaval nuclei (Fig. 1). These nuclei include the octaval nerve (VIII), the magnocellular octaval nucleus (MON), and the descending octaval nucleus (DON). Although not characterized in detail here, the more caudal brainstem also showed THir cell bodies and fibers in the area postrema, superior raphe, and vagal motor nuclei.

## Discussion

A comparative analysis of TH immunoreactivity (THir) between ancestral (surface) and derived (cave) forms of *Astyanax* revealed adaptive changes to the size and/or frequency of catecholaminergic neurons. Cavefish THir fibers appeared much more frequently than those of the surface fish, while THir cell body size and frequency varied between *Astyanax* by brain region. The cavefish has greater THir in the olfactory, tuberal, hypothalamic, and isthmic regions, while the surface fish has greater THir in the preoptic and ventral thalamic regions.

### TH neurons across teleostean fishes

THir neuron localization in the *Astyanax* brain is consistent with reports in other teleost fishes, once differences in anatomical nomenclature and boundary specification are taken into account. There is some variation in literature of ventral telencephalic nuclei reported as TH positive as well as fiber projections to the dorsal telencephalon. For example, THir neurons in the dorsal nucleus of the ventral telencephalon (Vd) appeared in surface *Astyanax*, zebrafish *Danio rerio* (Rink & Wullimann, 2000; Filippi et al., 2009), the Senegalese sole *Solea senegalensis* (Rodrıguez-Gomez et al., 2000), the Midshipman *Porichthys notatus* (Goebrecht et al., 2014), and the Elephant-nose fish *Gnathonemus petersii* (Meek & Joosten, 1993). THir Vd neurons were not present in the cave *Astyanax*, the Goldfish *Crassius auratus* (Hornby & Piekut, 1990), or the three-spined stickleback *Gasterosteus aculaeatus* (Ekström et al., 1990).

This discrepancy in THir ventral telencephalic nuclei and their projections is likely due to variations in the shape of the telencephalon across teleosts that maybe influenced by specializations to the olfactory system. The *Astyanax* cavefish has a shorter and broader ventral telencephalon compared to the surface fish and augmented olfactory detection capabilities (Hinaux et al., 2016). In the cavefish, olfactory tract (OT) projections from the ventral telencephalon to the dorsal region of the dorsal pallium (Dp) travel more laterally, while surface fish OT projections must travel more dorsally to reach Dp, in the vicinity of Vd boundaries.

Despite variations in telencephalic shape, THir in the olfactory bulb, basal telencephalon, preoptic, pretectal, tuberal, and isthmic regions is remarkably conserved across teleost fishes (see Rink & Wullimann, 2001), even with respect to larger type 2 neurons of the posterior tuberculum. Such neurons can be found in the brains of the Senegalese sole (Rodrıguez-Gomez et al., 2000), the Midshipman (Goebrecht et al., 2014), the three-spined stickleback (Ekström et al., 1990), the Goldfish (Hornby & Piekut, 1990), the brown trout *Salmo trutta fario* (Manso et al., 1993), the tench *Tinca tinca* (Brinon et al., 1998), the zebrafish *Danio rerio* (|Ma 1994a, 1994b, 1997, 2003; Rink & Wullimann, 2001; Filippi et al., 2009), and electric fish from both Gymnotiformes *Apternotus leptorrhyncus* (Sas et al., 1990) and Mormyridae *Gnathonemus petersii* (Meek & Joosten, 1993). For a more inclusive analysis of THir in bony fishes see Meek (1994).

The evolutionary conservation of THir nuclei across teleost fishes allows us to make predictions about catecholamine specification and connectivity in *Astyanax*. For example, comparative TH and Dopamine-β-hydroxylase immunohistochemistry studies in the zebrafish brain show no NA neurons rostral to the locus coeruleus (LC; Ma, 1997). As such, the THir nuclei in *Astyanax* that show marked similarities to those of the zebrafish are likely also dopaminergic. In addition, tract tracing studies performed in zebrafish and goldfish allow us to extrapolate similar connectivity in *Astyanax*. DiI injections into the zebrafish ventral pallium (Vv and Vd) doubled labeled Type 2 THir neurons of the posterior tuberculum (Rink & Wullimann, 2001), suggesting a similar ascending dopaminergic system from *Astyanax* Type 2 THir neurons.

In mammals, DA neurons modulate response behavior through ascending projections to the striatum that influence the motor and limbic pathways (Purves et al., 2014). Ascending DA modulation in teleost likely uses a similar mechanism, although identification of homologous structures is still ongoing. Zebrafish connectivity from type 2 neurons in the posterior tuberculum to the ventral pallium may indicate dopaminergic innervation of the “motor loop”, while the caudal hypothalamus (Hc) THir population may then serve as the dopaminergic modulation of the “limbic loop” (Rink & Wullimann, 2001; Rink & Wullimann, 2002). This hypothesis is consistent with our results as the cavefish is 1.5 times more active than surface fish, when regularly fed, and has larger and more numerous Type 2 neurons of the posterior tuberculum than the surface fish (Salin et al., 2010; Duboué et al., 2011). Although there were also more Hc THir cells of larger size in the cavefish, direct connections from these neurons to the telencephalon have yet to be reported in teleosts.

### Catecholaminergic cave adaptation

The same brain regions with greater cavefish THir are involved in the regulation of cave adaptive traits and behaviors. The lack of light influenced the enhancement of non-visual sensory systems, while the lack of both predation and nutrients likely drove the increased exploratory behaviors seen in the cavefish. We find greater THir in cavefish brain nuclei that process olfactory, gustatory, auditory and mechanosensory information as well as those that may serve as the dopaminergic innervation of motor circuitry. These results allow us to conclude not only that modifications in catecholamine neurons facilitated the evolution of cave adapted traits and behaviors, but that the increased exploratory behaviors of the cavefish are likely a product of finding limited food in the darkness.

#### Sensory cave adaptation

Early developmental degeneration of the cavefish eyes is offset by adaptations that enhance non-visual sensory modalities from those of the surface ancestor. An increase in cranial sensory neuromasts augments the mechanosensory system (Yoshizawa et al., 2012), while additional taste receptors line a broader jaw to better locate food (Yamamoto et al., 2009; Shiriagin & Korsching 2018), and odor detection sensitivity is enhanced by several orders of magnitude (Hinaux et al., 2016). The cavefish auditory system, surprisingly, appears to have no differences in sensitivity from the surface fish. However, the single investigation into *Astyanax* audition did reveal wider pure tone detection capacities than any other species tested at the time, implying that ancestral auditory characteristics may have already been sufficient for cave life (Popper, 1970).

The differences in THir in sensory processing nuclei reported between cave and surface *Astyanax* are therefore likely a reflection of DA potentiation of non-visual sensory systems in the cavefish. The surface fish showed greater THir in visual processing centers while the cavefish showed an increase in TH modulation of nuclei regulating not only olfaction, but also gustation, audition, and mechanosensation. The cavefish olfactory system promotes chemo-orientation at remote distances and initially induces food seeking behaviors (Kasumyan et al., 2013). This mostly involves running the mouth along the bottom towards the detected odor in back and forth patterns (Kowalko et al., 2013). The external taste buds along the bottom of the jaw then play an important role in successfully locating and eventually capturing food at more proximal distances (Schemmel 1967, 1974, 1980; Kasumyan & Marusov, 2018).

##### Olfaction

In the cavefish, modifications in sonic hedgehog signaling during development produced larger olfactory bulbs (OB) with more interneurons than in the surface fish (Menuet et al., 2007). Such an increase in OB interneurons has been positively correlated with enhanced odor discrimination (Scotto-Lomassese et al., 2003). To this end, we found a much greater quantity of THir OB cells in the cavefish. The cell bodies of these neurons were of smaller size compared to the surface fish and located on the border of the internal and external cell layers. The denser THir fiber projections from these neurons to the granule cell layer suggest a regulation of receptor cell input to mitral cells. This would enhance olfactory discrimination by increasing the lateral inhibition of mitral activity (for review see Kermen et al., 2013).

An increase in catecholaminergic neurons in the OB may also augment memory association to odorants. Studies in the zebrafish show projections from the ascending noradrenergic LC to the OB (Ma, 1994b), while studies in the carp support not only the presence of both dopamine and noradrenaline in the OB, but that long-term potentiation (LTP) following tetanic stimulation of the OT at the mitral-to-granule cell synapse is enhanced by noradrenergic signaling (Satou et al., 2006).

Further support for the link between increased catecholaminergic modulation and olfactory adaptation can be found in the asymmetric THir in the olfactory bulb of the Senegalese sole. Like most flatfish, the sole has one nostril on its blind side and an enlarged nostril on its sighted side. The OB of the enlarged nostril is also larger and displays an increase in THir cells and fibers compared to the blind nostril OB (Rodrıguez-Gomez et al., 2000). The differences between sole OBs closely resembles the variation between surface and cave *Astyanax* OB THir with the cavefish representing the enlarged OB.

Olfactory information is sent from the OB along mitral cell axons through the OTs for higher order processing in the forebrain (Von Bartheld et al., 1984). The Dp and Vv serve as the teleostean equivalents of the mammalian olfactory cortex and septal area, respectively. The Vv has reciprocal connections with the hypothalamus and both Vv and Dp nuclei are innervated by the noradrenergic LC and the dopaminergic posterior tubercle (Rink & Wullimannn, 2004; Schärer et al., 2012). The increased density of these fiber projections in the cavefish compared to the surface fish indicate a role of catecholamine modulation in the processing of olfactory information. The greater response strength of Vv neurons to odorant mixtures suggests a role in the mediation of the behavioral response to odorants in zebrafish (Yaksi et al., 2009). Action potentials in Dp were characterized by odor specific firing patterns triggered by simultaneous activity of converging mitral cell input and were strongly influenced by GABAergic inhibition (Yaksi et al., 2009), suggesting a role in the formation of odor memories.

Tract tracing studies in zebrafish also show a circuit connecting the OB to the tuberal region through the intermediate zone of the ventral pallium (Vi), which has been proposed as the teleostean correlate to the medial amygdala (Biechl et al., 2017). In mammals, the medial amygdala associates emotion with chemosensation and has functional subdivisions that mediate reproductive and defensive behaviors (Scalia & Winans, 1975; Lehman et al., 1980; Fernandez-Fewell and Meredith, 1994; Canteras et al., 1995; McGregor et al., 2004; Choi et al., 2005; Brennan & Zufall, 2006; LeDoux, 2012; Keshavarz et al., 2014). Vi THir was greater in the surface fish than the cavefish and may reflect the loss of defensive, alarm behaviors in adaptation to cave life (Pfeiffer, 1996; Hinaux et al., 2015). Intriguingly, DA was shown to selectively decrease inhibitory odor responses in Dp (Schärer et al., 2012), which, in the cavefish, could indicate further fine tuning within the circuit to facilitate memory association to particular odorants or even spatial odor mapping.

LOT projections through Dp also terminate in the habenula (Ha), which is highly conserved among teleosts and serves as a relay center from the telencephalon to the brainstem (Bianco & Wilson, 2008). Although tract tracing studies in the zebrafish brain show olfactory projections to the habenula in an asymmetric fashion (Miyasaka et al., 2009), we found very low immunoreactivity of fibers in this region that were only visible at magnifications greater than 20x and with no apparent asymmetries. Goldfish, a closer relative of *Astyanax* than zebrafish, also show little to no THir in the habenula (Hornby et al., 1987), despite its role in processing olfactory information.

##### Gustation

The cavefish feeds by running its enlarged jaw along the bottom in the proximity of a detected food source. The enhanced gustatory system spans the lower jaw and enables chemo detection of food particles in the benthic region (Boudriot & Reutter, 2001). Cavefish enhanced gustatory sensitivity arises from an increased number of tastebuds with additional sensory receptor cells, in addition to greater axon densities of some tastebud subtypes (Schemmel, 1967, 1974; Boudriot & Reutter, 2001 Varatharason et al., 2009). It was therefore not surprising to find greater TH innervation in higher order gustatory processing centers of the cavefish. The secondary gustatory nucleus (SGN) is somatotopically arranged and projects to the thalamic posterior nucleus (TPN) and the inferior lobe of the hypothalamus (IL; Wullimann, 1997; Rink & Wullimann, 1998; Folgueira et al., 2003). All three regions showed much greater THir in the cavefish than in the surface fish, although THir was predominantly in the caudal region of the TPN. Studies in mammals show that DA release is stimulated by palatable taste sensation and that feeding behaviors cease upon the addition of DA antagonists (Ahn & Phillips 1999; Hajnal and Norgren 2001; for review see Katz & Sadacca, 2011). Because taste is so strongly linked with reward, it is likely that THir in gustatory nuclei of the cavefish is increased dopaminergic signaling to enforce consumption of much needed nutrients.

##### Audition and mechanosensation

The increased TH modulation of the cavefish auditory nuclei, such as the octaval nerve, was unexpected, but may be a product of differences between cave and surface aquatic environment sound profiles. Other fish, such as the ancient Amblyopsids do show differences in hearing profiles between cave and surface species. High-frequency hearing loss in *Amblyopsid* cavefishes has been attributed to high intensity 1000 Hz peaks in background noise produced through cave amplification of water droplets and flowing streams (Niemiller et al., 2013). Despite this potential cave echo effect, the evolutionarily conserved ability of *Astyanax* cavefish to detect high pitch noises may be adaptive. In the Pachón cave, water inflow is less frequent than in the *Amblyopsid* cave system and brings much needed food to the nutrient depleted pool. As such, the sounds associated with flowing water during the rainy season would indicate an approaching food source and proceed even the olfactory system in the detection of food stimuli. Dopamine signaling to auditory processing centers, such as the octaval nucleus, would therefore associate roaring cave echoes with reward and initiate food seeking behaviors.

Sound and vibration are a mixed stimulus in water and auditory and mechanosensory systems send parallel projections to the same higher order nuclei (Finger & Tong, 1984; Echteler, 1985). Because cavefish have such a strong vibration attraction behavior (VAB; Yoshizawa et al., 2010), it is likely the THir of these sensory processing nuclei are, at least in part, the dopaminergic modulation of the lateral line system. Due to the increased detection capacities of cranial neuromasts and strength of cavefish VAB (Yoshizawa et al., 2010, 2011), we were surprised to not see THir in the anterior lateral line (ALL) nucleus. However, the ALL nucleus, along with the posterior lateral line nerves, sends afferents to the nearby medial octaval nucleus (MON) and descending octaval nucleus (DON), both regions with greater THir in the cavefish. This trend continues to the following level of auditory and mechanosensory processing, the torus semicircularis (TS), where the cavefish continue to show much great THir (for review see Wullimann, 1998). From here, lateral line information is sent to the dorsal region of the dorsal pallium (Dd; often combined with the medial region of the dorsal pallium in tract tracing studies in other teleosts; Echteler, 1985), which only shows THir in the surface fish. Unlike the cavefish, the surface fish has aversion behaviors to vibration stimuli, likely a product of predation pressures not found in the cave system. Surface fish Dd THir could, therefore, represent noradrenergic fear association to mechanosensory stimuli that has been lost in the cavefish. At the top trophic level, cavefish would experience no negative consequences from investigating vibration stimuli. As such, THir in the cavefish MON, DON, and TS is most likely a continuation of dopaminergic modulation of feeding behavior. Reward association with vibration stimulus would motivate investigation of a crucial potential food source rather than the fear association of the ancestral surface fish that faces greater survival threat from predation than starvation.

##### Vision

Adult cavefish have a degenerated eye, that is sunken into the orbit and covered by fat and skin (for review see Keene et al., 2015). It is commonly believed that they are completely blind (Romero et al., 2003). Surprisingly, despite their failure to respond to visual cues, the cavefish showed similar, albeit reduced, catecholaminergic modulation of visual nuclei as seen in the surface fish. This could be because visual regions receive enough innervation from the cavefish eye before it degenerates to induce their development or more likely, due to the incomplete allopatric speciation in *Astyanax*. In terms of evolutionary time, the cavefish is relatively young, with some divergence estimates as recent as the last 10,000 years (Fumey et al., 2018). As such, degeneration of a non-functional visual system may be ongoing and the catecholaminergic modulation of visual processing centers merely vestigial.

Tract tracing studies in bluegill (Northcutt & Butler, 1991), the three-spined stickleback (Ekström, 1984) show retinal termination fields in the ventral medial (VM) and ventral lateral (VL) thalamus as well as the suprachiasmatic nucleus (SCN). Although retinofugal projections to PPv are less consistent between species, PPv TH fibers extend ventro laterally along the base of the tectum where retinal termination fields were also reported (for review see Northcutt & Wullimann, 1988). Somewhat less surprisingly, THir was significantly reduced in the blind cavefish in these visual regions than in the surface fish. Surface fish THir cell bodies in the nuclei were larger and more numerous. PPv was the only exception to this where there was little difference between *Astyanax* forms. This may be because although it modulates centers that receive visual information, it may not receive visual cues itself and so is less affected by disparities in visual innervation in the cavefish.

The same nuclei that receive visual input were also shown to receive projections from the pineal gland (Ekström, 1984), which may explain their THir in the cavefish. Although cavefish in the Pachón cave are never exposed to light, those raised in the laboratory receive regular light stimulus from the time of inception. Although blind, the cavefish is also mostly transparent and may receive light stimulus directly to the conserved pineal organ through the skull (Grunewald-Lowenstein, 1956; Omura, 1975; Herwig, 1976; Yoshizawa & Jeffery, 2008). Reports in zebrafish describe not just the pineal gland, but the entire brain as light sensitive, although the relevant opsins are still being identified (Moore & Whitmore, 2014). This is of particular relevance because the pineal gland plays a much greater role in the regulation of the circadian clock in teleosts than in mammals (for review see Ben-Moshe et al., 2014). Disparages in the oscillation of circadian clock genes have been reported between *Astyanax* raised in the lab that follow a 12-hour light/dark cycle and those native to the cave environment that do not experience oscillations (Beale et al., 2013). Light input to the pineal gland in lab raised cave *Astyanax* may, therefore, induce clock gene expression and the catecholaminergic modulation of nuclei that process light stimulus in the blind cavefish.

Retinal and visual nuclei afferents eventually all converge on optic tectum (TOp). The TOp is the main visual processing center of the teleost brain and is arranged in layers with a loose resemblance to the mammalian cortex. Although not quantitatively analyzed, TH fibers were present within the TOp of both *Astyanax* fish, despite the smaller size and decreased visual innervation of the cavefish tectum (Fig. 4; Soares et al., 2004). Electrophysiological recordings from the cavefish TOp revealed that somatic innervation replaces the spatial mapping ordinarily created by the visual system, with somatic input from the head of the cavefish taking up a larger area than the rest of the body (Voneida & Fish, 1984). In accordance with behavior observations of cavefish placed in novel aquaria, somatosensory signaling to the TOp likely allows the cavefish to map its surroundings by bumping into objects with its head (Gertychowa, 1970). Because the somatosensory system compensates for the role normally played by the visual system, the TOp is likely innervated by catecholamines in a similar manner in both *Astyanax* forms.

#### Homeostatic cave adaptation

Cavefish catecholamine adaptation to the nutrient poor cave environment not only augmented traits to acquire food, but also those to conserve energy. The consistency of the cave environment for most of the year decreases the need to secrete hormones, moderate body temperature, and follow a 24-hour circadian rhythm. Summertime in the Sierra del Abra region of Mexico can raise the sunlit surface waters up to 28.1°C (Simon et al., 2017), while the cave waters remain at a relatively constant mean temperature of 23.1°C all year long (Elliot, 2015). In fact, the only rise in cave temperature occurs when the spring rains bring the warmer surface waters flooding into the stagnant cave system. This event triggers reproductive behaviors in the cave fish, likely due to temperature effects on hormone secretion and the influx of dissolved nutrients from the surface. Because homeostasis is maintained by the external environment, cavefish are likely able to stretch limited energy resources between rare feeding opportunities. In support of this theory, we found that differences in catecholamine neurons between surface and cavefish reflected differences in the regulation of energy homeostasis.

##### Endocrine Secretion

DA projections from the anterior parvocellular preoptic nucleus (PPa) terminate in the pituitary where they regulate the secretion of hormones such as gonadotropin, growth hormone (GH), and α-melanin-stimulating hormone (α-MSH; Gordon, 1958; Arosio et al., 1980; Wong et al., 1992,1993; Fontaine et al., 2015). PPa THir neurons had a larger diameter in surface fish, although the cavefish had many more neurons of smaller size. These differences likely reflect environmentally dependent requirements for the inhibition of endocrine release and the energy demands for endocrine secretion.

In fishes, warm water is required for gonadotropin release (Gordon, 1958; Umminger 1978), which may explain why cavefish only reproduce following warm water floods. Further in zebrafish, DA has an inhibitory effect on the secretion of gonadotropin (Fontaine et al., 2015). The natural cooling of receding flood water in the cave system leaves the cavefish with little need for prolonged DA regulation of gonadotropin release. The surface fish, however, spends the summer months in much warmer water and the larger PPa cells may be necessary to inhibit the release of reproductive hormone following spring spawning events.

Although the need to regulate gonadotropin release is likely diminished in cavefish, the greater number of smaller PPa neurons suggests lower level continual inhibition of GH and α-MSH secretion. Unlike most fishes, Pachón cavefish experience determinate growth (Simon et al., 2017). This is presumably to limit the greater energy demands of a larger body size in a nutrient poor environment. Although the mechanisms that halt cavefish growth at a genetically predetermined size have yet to be investigated, dopaminergic inhibition of GH release may play a key role in restricting increases in body size that are not sustainable by the limited cave resources.

Starvation adaptation has also resulted in a satiation resistant cave phenotype (Strickler & Soares, 2011). α-MSH secretion inhibits feeding behaviors in rainbow trout (Schjolden et al., 2009) and goldfish (Cerdá-Reverter et al., 2003) and increased inhibition of its release in cavefish may contribute to their insatiability. As such, PPa DA inhibition of GH and α-MSH secretion would provide adaptive advantages to survival on limited food reserves.

##### Temperature regulation

While PPa catecholamine release inhibits endocrine release, posterior parvocellular preoptic nucleus (PPp) catecholamine signaling induces homeostatic adjustments to changes in water temperature. Intracranial injections of L-dopa into the Mediterranean chromis, *Chromis chromis*, resulted in an increase in body temperature of the fish (Green & Lomax, 1976). Similarly, intracranial injections of DA and NA into what we presume to be the PPp of the Goldfish induced behavioral selection towards colder water (Wollmuth & Crawshaw, 1989). This preference for cold water likely reflects the increase in body temperature modulation via increases in catecholamine signaling in PPp. The larger size and quantity of PPp cell bodies in the surface fish as opposed to the cavefish, may even reflect a preference of the ancestral state for colder water that could have facilitated entry into the cave system. While the derived decrease in cavefish PPp THir may reflect the consistency of Pachón cave water temperature and even an energy conservation mechanism.

##### Sleep as energy conservation

The absence of light stimulus has a pronounced effect on the sleep/wake cycles of the cavefish. Not only are they completely released from visual zeitgebers and internal circadian rhythms, they are completely food driven (Beale et al., 2013; Jaggard et al., 2017, 2018). Pachón cavefish sleep more when starved and less when fed regularly. In fact, regular feeding dramatically decreases cavefish sleep intervals and duration (Jaggard et al., 2017). Most starving animals increase their survival probability by forgoing sleep in favor of foraging to use their remaining energy reserves to seek needed nutrients (McDonald & Keene, 2010). Cavefish, however, have nothing to be gained by continuously searching the barren confines of their habitat. Instead these animals sleep to conserve energy until a possible feeding opportunity arises. As a result of their enhanced non-visual sensory capabilities, cavefish have a dramatically reduced arousal threshold to that allows them to awake with each novel disturbance (Jaggard et al., 2017)

The LC is one of the three major brain nuclei responsible for promoting the wake state, through noradrenergic signaling (for review see Saper et al., 2005). The prolonged wakefulness of the cavefish is correlated in the greater number and size of THir LC neurons compared to the surface fish. This additional noradrenergic signaling likely alerts the cavefish to novel stimuli and promotes the wake state while the stimulus is investigated.

### Conclusions

Here we report a correlation between changes in the size and/or number of THir neurons in *Astyanax* and adaptations to cave life. Catecholamine signaling was more pronounced in the surface fish visual system and with regards to temperature dependent homeostasis. Catecholamine signaling in the blind cavefish was more pronounced in non-visual sensory systems, regions that regulate attention and locomotor activity, and the inhibition of hormones that regulate growth and satiety. These increases in catecholaminergic neuron size and frequency likely allow the cavefish to make the most of every rare feeding opportunity by quickly detecting and locating food in absolute darkness, while preserving energy when food is scarce. As such, evolutionary changes in catecholamine neurons may underlie the emergence of new adaptive behaviors.

## Acknowledgements

We would like to thank Dr. Kristen Severi for her helpful comments on the manuscript and Liam Dishy for help with anatomical measurements.

Grant support: NIH R15EY027112

The data that support the findings of this study are available from the corresponding author upon reasonable request.

